# Estrogen regulation and functional role of FGFR4 in estrogen receptor positive breast cancer

**DOI:** 10.1101/2024.03.18.585626

**Authors:** Kai Ding, Lyuqin Chen, Kevin Levine, Matthew Sikora, Nilgun Tasdemir, David Dabbs, Rachel Jankowitz, Rachel Hazan, Osama S Shah, Jennifer M Atkinson, Adrian V Lee, Steffi Oesterreich

## Abstract

**Background:** Resistance to endocrine therapy is a major challenge of managing estrogen receptor positive (ER+) breast cancer. We previously reported frequent overexpression of FGFR4 in endocrine resistant cell lines and breast cancers that recurred and metastasized following endocrine therapy, suggesting FGFR4 as a potential driver of endocrine resistance. In this study, we investigated the role of FGFR4 in mediating endocrine resistance and explored the therapeutic potential of targeting FGFR4 in advanced breast cancer.

**Methods:** A gene expression signature of FGFR4 activity was examined in ER+ breast cancer pre- and post-neoadjuvant endocrine therapy and the association between FGFR4 expression and patient survival was examined. A correlation analysis was used to uncover potential regulators of FGFR4 overexpression. To investigate if FGFR4 is necessary to drive endocrine resistance, we tested response to FGFR4 inhibition in long term estrogen deprived (LTED) cells and their paired parental cells. Doxycycline inducible FGFR4 overexpression and knockdown cell models were generated to examine if FGFR4 was sufficient to confer endocrine resistance. Finally, we examined response to FGFR4 monotherapy or combination therapy with fulvestrant in breast cancer cell lines to explore the potential of FGFR4 targeted therapy for advanced breast cancer and assessed the importance of PAM50 subtype in response to FGFR4 inhibition.

**Results:** A FGFR4 activity gene signature was significantly upregulated post neoadjuvant aromatase inhibitor treatment, and high FGFR4 expression predicted poorer survival in patients with ER+ breast cancer. Gene expression association analysis using TCGA, METABRIC and SCAN-B datasets uncovered ER as the most significant gene negatively correlated with FGFR4 expression. ER negatively regulates FGFR4 expression at both the mRNA and protein level across multiple ER+ breast cancer cell lines. Despite robust overexpression of FGFR4, LTED cells did not show enhanced responses to FGFR4 inhibition compared to parental cells. Similarly, FGFR4 overexpression, knockdown or hotspot mutations did not significantly alter response to endocrine treatment in ER+ cell lines, nor did FGFR4 and fulvestrant combination treatment show synergistic effects. The HER2-like subtype of breast cancer showed elevated expression of FGFR4 and an increased response to FGFR4 inhibition relative to other breast cancer subtypes.

**Conclusions:** Despite ER-mediated upregulation of FGFR4 post endocrine therapy, our study does not support a general role of FGFR4 in mediating endocrine resistance in ER+ breast cancer. Our data suggests that specific genomic backgrounds such as HER2 expression may be required for FGFR4 function in breast cancer and should be further explored.

## Introduction

Seventy percent of breast cancers are estrogen receptor positive (ER+) and are treated with endocrine therapy that antagonizes estrogen receptor signaling [1, 2]. Endocrine therapy has revolutionized the treatment of ER+ breast cancer by reducing the risk of 10-year recurrence and the risk of death by 50% and 30% respectively [3]. However, ∼30% of patients with ER+ breast cancer will develop recurrence which is refractory to endocrine therapy [4].

Multiple mechanisms driving endocrine resistance in breast cancer have been reported, including deregulation of the ER pathway via 1) ESR1 mutations, fusions or loss of expression, 2) increased activation of receptor tyrosine kinase pathways including ERBB2, EGFR, FGFR1, FGFR2, and IGF1-R, 3) alteration of PI3K and MAPK pathways, 4) deregulation of transcriptional regulators including MYC, FOXA1 and ARID1A, and 5) tumor metabolic reprogramming [5–7]. Despite the successful development of combination therapies targeting ER and CDK4/6, mTOR or receptor tyrosine kinases, endocrine resistance remains a significant clinical challenge.

In search of drivers of endocrine resistance, we previously performed RNA sequencing (RNAseq) analysis of 12 long term estrogen deprived (LTED) breast cancer cell lines and their paired endocrine sensitive parental cells and identified the consistent overexpression of FGFR4 [8]. Furthermore, we and others have reported frequent overexpression of FGFR4 in breast cancer metastases post endocrine therapy compared to paired primary tumors, especially in invasive lobular breast carcinoma (ILC) [8, 9]. ILC has been reported to provide a permissive background enhancing growth factor receptor signaling and function [10, 11]. Further, high FGFR4 expression is associated with an inferior clinical benefit of tamoxifen treatment [12]. Notably, no consistent upregulation of FGFR1-3 was observed post endocrine therapy, nor an association with outcome with tamoxifen treatment [12, 13]. Together, these data strongly suggest FGFR4 as a potential driver of endocrine resistance in breast cancer.

FGFR4, along with FGFR1-3, belongs to the fibroblast growth factor receptor family, a highly conserved receptor tyrosine kinase family [14]. FGFR4 shares the least homology compared to other FGFRs [15]. Notably, FGFR4 has a unique residue C552 within the kinase domain enabling the design of FGFR4 specific inhibitors. Several FGFR4 specific inhibitors have been tested in recent clinical trials for hepatocellular carcinoma, including Fisogatinib (BLU-554) [16], H3B-6527 [17], and INCB062079 [18] (recently reviewed in [19]). Like other FGFRs, canonical FGFR4 signaling is dependent upon the binding of FGF ligands enabled by heparin or klotho proteins, which leads to the downstream activation of phospholipase γ (PLC γ), FGFR substrate 2 (FRS2), the signal transducer and activator of transcription (STAT) signaling pathway, Wnt/GSK-3 β /β-catenin pathway, and Macrophage Stimulating 1 or 2 (MST1/2) [13]. FGFR4 shares most of the FGF ligands with other FGFRs including FGF1/2/4/6/8/17, with FGF19 reported to predominately bind to FGFR4 in concert with beta klotho [20]. Noncanonically, FGFR4 has been shown to function through interaction with other genes in a ligand independent manner, including NCAM [21], N-cadherin [22], EphA4 and ephexin1 [23], and MT1-MMP1 [13, 24].

Aberrant activation of FGFR4 signaling through receptor or ligand overexpression, FGFR4 mutation, or a G388R single nucleotide polymorphism (rs351855) has been reported to promote progression and mediate treatment resistance in multiple types of cancer [13]. The oncogenic functions of FGFR4 are well investigated in hepatocellular carcinoma [25–27] and rhabdomyosarcoma (RMS) [28, 29], where FGFR4 is reported to be the major driver of tumor progression, and in these cancer types, inhibition of FGFR4 can impede tumor growth both *in vitro* and *in vivo*. In breast cancer, FGFR4 has been reported to be involved in tumor oncogenesis [30], maintaining the survival of breast cancer cell lines co-expressing FGFR4 and FGF19 [31], and mediating the resistance to chemotherapy [32]. Mutation of FGFR4 is relatively rare in cancer, with metastatic melanoma showing the highest mutation rate of 5.5% [8]. Breast cancer is the only cancer type exhibiting a significant enrichment of FGFR4 mutation in metastases (2.2%) compared to the primary tumor (0.45%), suggesting a role in therapy resistance and metastasis. Two FGFR4 hotspot mutations, the N535D/K and V550E/L/M, were previously identified and investigated in RMS, where they were shown to promote tumor proliferation and metastasis in xenograft models through enhanced FGFR4 autophosphorylation [29, 33]. These two hotspot mutations have also been shown to mediate resistance to FGFR4-specific inhibitors and importantly, we have previously reported the almost exclusive enrichment of these hotspot mutations in breast cancer metastases compared to other tumor types in adults [8].

Despite strong evidence supporting FGFR4 as an important driver and mediator of endocrine resistance in breast cancer, FGFR4 is understudied compared to other FGFRs likely due to its lower frequency of somatic genomic alterations. In this study, we investigated the mechanism of FGFR4 overexpression post endocrine therapy, the role of FGFR4 in mediating response to endocrine therapy, and the therapeutic potential of targeting FGFR4 in advanced breast cancer.

## Results

### A FGFR4 activity gene signature is upregulated in response to endocrine therapy and high FGFR4 expression predicts worse patient survival in ER+ breast cancer

To examine if FGFR4 is activated post endocrine therapy, we calculated the GSVA score of a FGFR4 activation signature previously developed by Garcia-Recio S et al. [34] using RNA sequencing (RNAseq) data of paired estrogen receptor positive (ER+) primary breast cancer pre-and post-90-day neoadjuvant aromatase inhibitor (AI) therapy [35, 36]. Although FGFR4 mRNA itself was not upregulated (Figure S1), the FGFR4 signature was significantly elevated post AI treatment (Figure 1A). To investigate if FGFR4 was associated with ER+ breast cancer survival, we quantified FGFR4 expression in 123 ER+ ILC samples from patients with available data on distant recurrence-free survival (supplementary table 1). Patients with ILC with high FGFR4 expression had significantly worse distant recurrence free survival than patients with low FGFR4 expressing ILC (Figure 1B). We expanded this study to ER+ breast cancer from the METABRIC cohort (n=1505), including both ILC and invasive ductal breast cancer (IDC) (breast cancer of no special type, NST). High FGFR4 expression was again associated with worse disease-free survival, suggesting that enhanced FGFR4 expression might be associated with a poorer response to endocrine therapy and therefore worse survival of patients with ER+ disease (Figure 1C).

**Figure 1.**
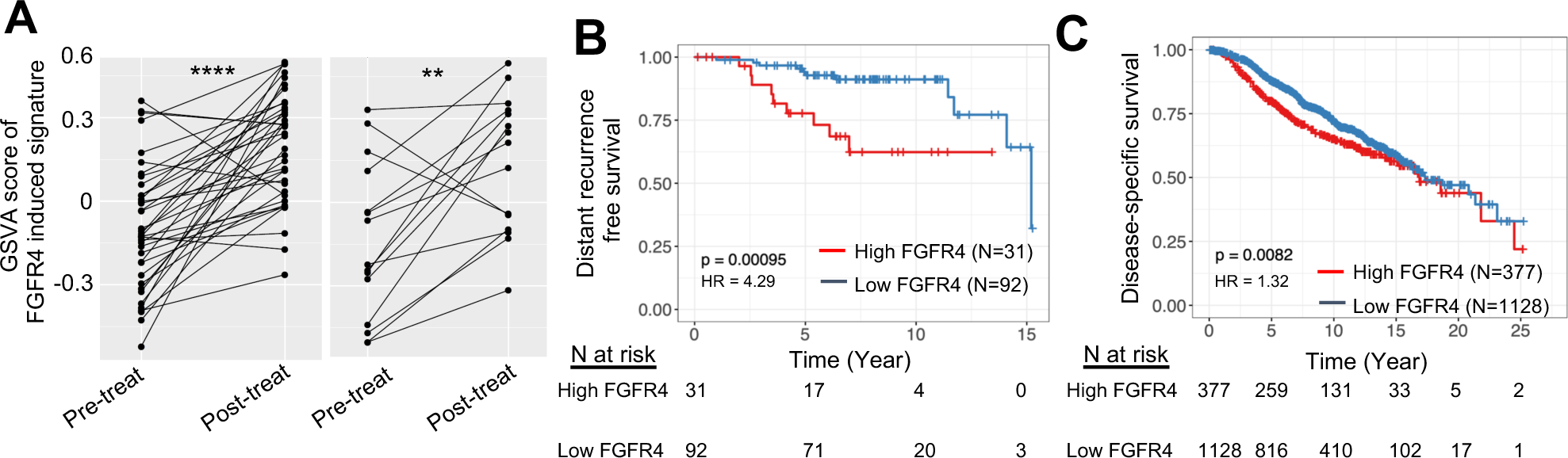
Clinical evidence supports a potential role of FGFR4 in mediating endocrine resistance. (A) GSVA score of FGFR4 induced signature pre and post 90 days of neoadjuvant aromatase inhibitor treatment in ER+ breast cancer using data from GSE59515+GSE5537 (left) and dbGaP (phs000472) (right). p value was calculated by paired Wilcoxon test. (B) Association between FGFR4 expression levels with distant free survival in WCRC cohort and (C) with disease specific survival in METABRIC cohort. ER+ breast cancer from both cohorts were included for the analysis and the third quantile (Q3) was used as cutoff defining FGFR4 high and FGFR4 low group. p value was calculated using log rank test and hazard ratio was calculated by Cox proportional hazard regression analysis. * p < 0.05, ** p < 0.01, *** p < 0.001, **** p < 0.0001.

### FGFR4 expression is negatively regulated by ER

To understand the mechanisms of *FGFR4* overexpression, we set out to determine which genes were correlated with FGFR4 expression in ER+ primary tumors. Using TCGA data, we identified 28 proteins (or post-translationally modified proteins) that were significantly positively correlated with FGFR4 mRNA, while 26 proteins were significantly negatively correlated with FGFR4 mRNA (Figure 2A). The strongest positive associations were seen with *HER2*_pY1248 and *EGFR*_pY1068 while progesterone receptor (PR), ERALPHA (ER) and ERALPHA_pS118 were negatively associated with FGFR4 mRNA. Considering the critical role of ER as master transcriptional regulator and drug target in ER+ breast cancer, we further examined the correlation between FGFR4 mRNA and ESR1 mRNA expression in ER+ samples from TCGA (n=808), METABRIC (n=1505) and SCAN-B (n=2945) RNAseq data sets. Consistently, FGFR4 was significantly negatively correlated with ESR1 expression in all three data sets (Figure 2B), suggesting that ER may act as a negative regulator of FGFR4 transcription. Two ER binding peaks were identified 63kb upstream and 22kb downstream of FGFR4 transcription start site in MCF7, ZR75-1, T47D and MDA-MB-134 cell lines upon estradiol (E2) treatment (Figure S2). To experimentally test whether ER is a negative regulator of FGFR4, FGFR4 mRNA and protein expression were examined in a panel of ER+ breast cancer cell lines with or without ESR1 siRNA knockdown. These studies confirmed that FGFR4 mRNA (Figure 2C) and protein (Figure 2D) expression were upregulated upon ESR1 knockdown in all cell lines tested with the exception of MCF7 and SUM44PE. In support of this data, we further observed that estradiol (E2) treatment could induce downregulation of FGFR4 mRNA which was rescued by fulvestrant (ICI 182780) treatment (Figure 2E), again with the exception of MCF7 and SUM44PE cells. In summary, FGFR4 expression is negatively associated with ER expression in ER+ tumors, at least in part due to ER repressing transcription of FGFR4 mRNA.

**Figure 2.**
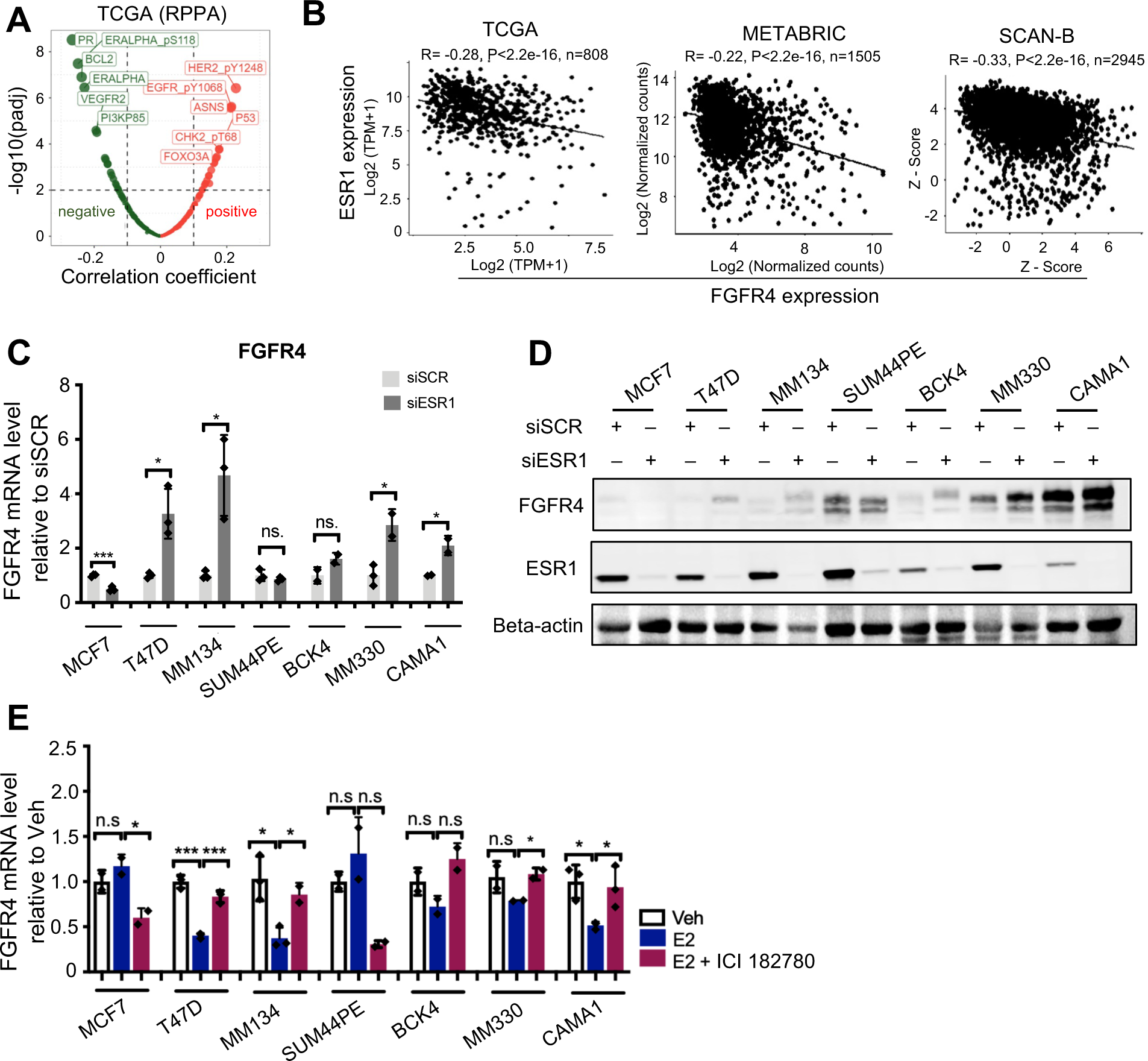
FGFR4 is negatively regulated by ER. (A) Spearman correlation analysis between FGFR4 mRNA expression and protein expression from TCGA RPPA data. (B) FGFR4 and ESR1 mRNA expression spearman correlation in ER+ samples of TCGA, METABRIC and SCAN-B data sets. R. spearman correlation coefficiency. (C) FGFR4 mRNA expression in ER+ cell lines treated with control (siSCR) or siESR1, quantified by qRT-PCR. Data was normalized to siSCR group for each cell line. Representative data from N=3 independent repeats. (D) FGFR4 protein expression in ER+ cell lines treated with control (siSCR) or siESR1, quantified by western blot. Representative data from N=3 independent repeats (E) FGFR4 mRNA expression in ER+ cell lines deprived in charcoal stripped serum (CSS) for 3 days and treated with Veh. (Vehicle), 1 nM estradiol (E2) or 1 nM E2 + 100 nM ICI 182780 for 24 hours. Data was normalized to Veh. treated group of each cell line. Representative data from N=3 independent repeats Error bar represents standard deviation (SD). p values for C, D and E are calculated by t test. * p < 0.05, ** p < 0.01, *** p < 0.001, **** p < 0.0001. n.s (nonsignificant)

### Despite increased FGFR4 expression, LTED models do not show an enhanced dependence on FGFR4 for survival, relative to parental cells

We have previously generated [37] multiple clones of long-term estrogen deprived (LTED) cells derived from ER+ MDA-MB-134 and SUM44PE cells, which grow independent of E2 and show consistent overexpression of FGFR4 (Figure S3A) compared to paired parental cells and can grow independent of E2. To examine if FGFR4 overexpression is required for the E2 independence of LTED cells, we generated doxycycline (Dox) inducible shRNA-mediated FGFR4 knockdown models in MDA-MB-134 and SUM44PE parental and LTED cells (Figure 3A, 3B). The growth of MDA-MB-134 (clones LB and LE) and SUM44PE (clone LA) LTED cells in CSS was only weakly inhibited by FGFR4 knockdown (Figure 3A, 3B). We also examined the response of LTED cells to FGFR4-inhibitors H3B-6527 and BLU554. MDA-MB-453, a positive control cell line bearing an activating FGFR4 mutation (Y367C), showed robust responses to H3B-6527 and BLU554, while MDA-MB-134 parental and LTED cells showed little response to either drug in 2D growth or colony formation assays (Figure 3C, 3D, 3E, 3F). SUM44PE cells exhibited stronger responses to H3B-6527 and BLU554 compared to MM134 cells, but there was no consistent significant difference in response between SUM44PE parental and LTED cells (Figure 3G, 3H, S3B, S3C). We further assessed response to H3B-6527 in a pair of MCF7 parental and LTED cells [38], which also failed to reveal a difference in sensitivity between parental and LTED cells despite the upregulation of FGFR4 in the LTED models (Figure S3D-E).

**Figure 3.**
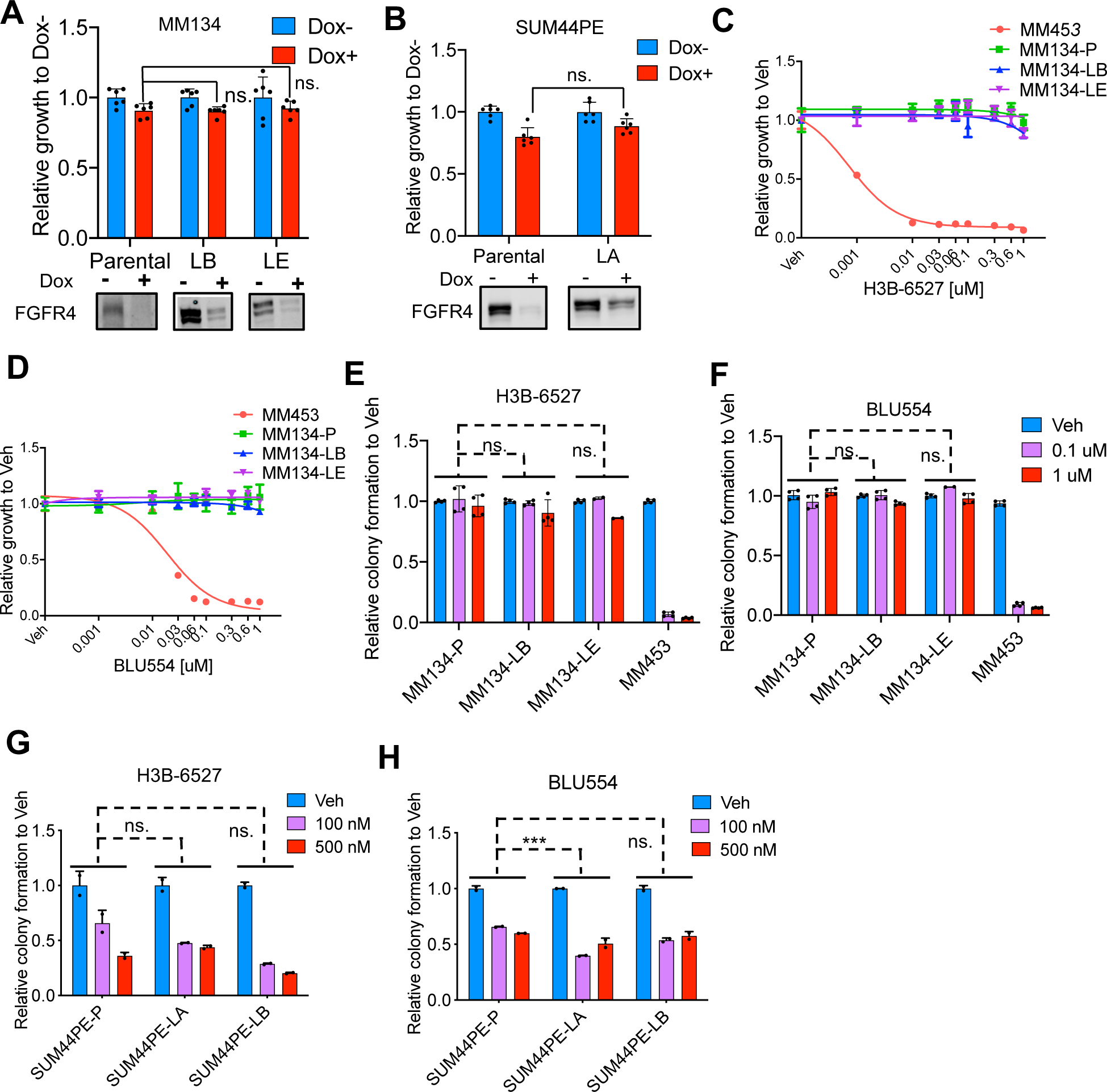
Despite increased FGFR4 expression, LTED models do not show an enhanced dependence on FGFR4 for survival, relative to parental cells. (A) Relative growth of MDA-MB-134 parental, LTED-B (LB) and LTED-E (LE) cells and (B) SUM44PE parental and LTED-A (LA) cells in CSS with doxycycline (Dox) inducible knockdown of FGFR4. FGFR4 knockdown was confirmed by western blot. Cell number was quantified by FluoReporter Assay. Data was normalized to Dox-group of each cell line. Representative data from N=2 independent repeats for (A), N=3 for (B). (C) Growth inhibition of MDA-MB-134 P (Parental), LB, LE and MDA-MB-453 cells in response to H3B-6527 and (D) BLU554 treatment for 7 days. Cell number was quantified by FluoReporter Assay. Data was normalized to Veh. treated group of each cell line. Dose response curve was simulated using a nonlinear regression model with variable slopes by Graphpad PRISM. Representative data from N=3 independent repeats for (C) and (D). (E) Colony formation of MDA-MB-134 cells in response to H3B-6527 and (F) BLU554 treatment, and colony formation of SUM44PE cells in response to (G) H3B-6527 and (H) BLU554 treatment for 2 weeks. Colonies were quantified by crystal violet staining. Data was normalized to Veh. treated group of each cell line. Representative data from N=3 independent repeats for E, G and H and N=2 for F. All error bars represent standard deviation. p values for A and B were calculated by t test, and for E-H were calculated with two-way ANOVA. * p < 0.05, ** p < 0.01, *** p < 0.001, **** p < 0.0001. ns. (nonsignificant)

### FGFR4 expression alteration does not alter response to endocrine therapy in ER+ breast cancer models

To investigate whether FGFR4 overexpression confers resistance to endocrine therapy, we generated Dox-inducible FGFR4 overexpression models using four ER+ cell lines with low endogenous expression of FGFR4, including MDA-MB-134, MCF7, T47D, and BCK4 cells (FigureS4A). Cell growth in CSS to mimic estrogen depletion in the presence or absence of Dox (and thus presence or absence of FGFR4) was measured using 2D growth assay. No significant differences in cell growth in the presence or absence of Dox were observed, suggesting that FGFR4 does not contribute to cell survival in this acute estrogen depleted setting (Figure 4A, S4B). Further, FGFR4 overexpression did not alter response to ICI 182780 in MDA-MB-134, MCF7 orT47D cells (Figure S4C-E).

**Figure 4.**
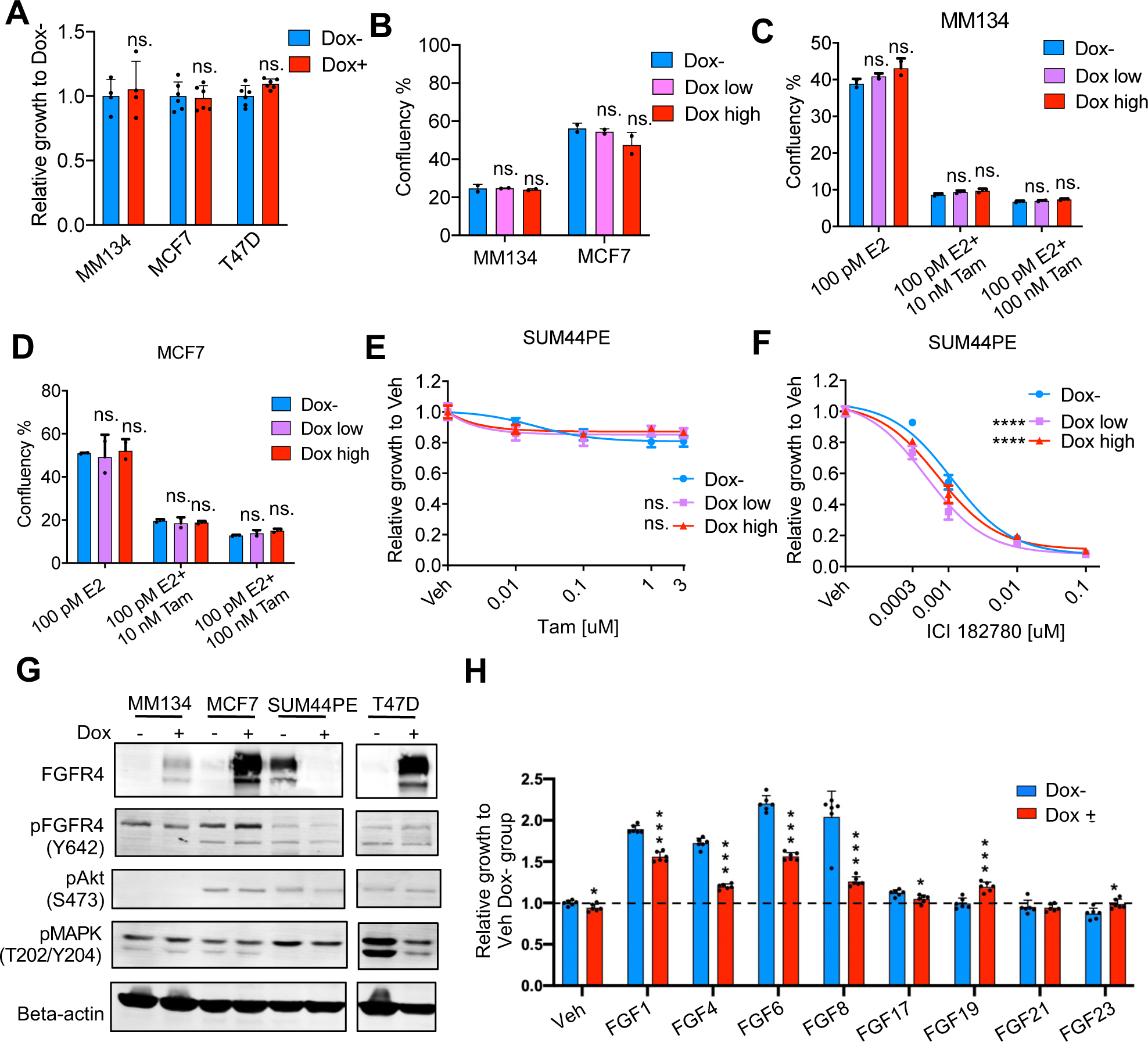
FGFR4 expression alteration did not alter response to endocrine therapy in models tested. (A) Relative growth of MDA-MB-134, MCF7 and T47D cells in CSS with and without Dox induced overexpression of FGFR4. Cell numbers were quantified using FluoReporter assay after 7 days. Data was normalized to Dox-group of each cell line. Representative data from N=2 independent repeats. (B) Confluency of MDA-MB-134 and MCF7 cells in CSS with and without Dox induced overexpression of FGFR4. Cell confluency was quantified using Incucyte live imaging system after one month of culture. Representative data from N=3 independent repeats. (C) Confluency of MDA-MB-134 and (D) MCF7 cells in E2 supplemented CSS in the presence and absence of Tam (Tamoxifen) treatment with and without Dox induced overexpression of FGFR4. Cell confluency was quantified using Incucyte live imaging system after one month of culture. Representative data from N=3 independent repeats. (E) SUM44PE dose responses to Tam and (F) ICI 182780 with and without Dox induced knockdown of FGFR4. Cell numbers were quantified using FluoReporter assay after 7 days. Data was normalized to Veh. treated group of each condition. Response curves were simulated using nonlinear regression model with variable slopes by PRISM. Representative data from N=2 independent repeats. (G) Western blot assessment of FGFR4 and downstream signaling levels in cell models with and without Dox induced overexpression (MM134, MCF7, T47D) or knockdown (SUM44PE) of FGRF4 for 48 hours. Representative data from N=2 independent repeats. (H) T47D cell growth in CSS with and without Dox induced overexpression of FGFR4 in the presence and absence of FGFs (20 ng/ml for FGF1/4/6/8/17, 100ng/ml for FGF17/19/21/23). Viable cell numbers were quantified using PrestoBlue assay after 7 days. Data was normalized to Veh. treated Dox-group. Representative data from N=3 independent repeats. Error bar in all figures represent standard deviation. p values comparing Dox+ group to Dox-group for figures A, B, C, D and H were calculated using t test, and for figures E and F were calculated using two-way ANOVA. * p < 0.05, ** p < 0.01, *** p < 0.001, **** p < 0.0001. ns. ( nonsignificant)

To examine if long-term FGFR4 overexpression was required to mediate endocrine resistance, MDA-MB134 and MCF7 cells were cultured in CSS (Figure 4B) or CSS supplemented with E2, in the presence or absence of Tamoxifen (Tam) (Figure 4C, 4D) and treated with or without Dox for one month. Consistent with the prior data, this assay also revealed no significant differences between groups with and without FGFR4 overexpression. To corroborate this finding, we examined if FGFR4 knockdown can increase response to endocrine therapy in a panel of ER+ cells with high endogenous FGFR4 expression. SUM44PE cells did not show notable differences in response to Tam (Figure 4E) or ICI 182780 (Figure 4F) treatment between cells with or without Dox induced shRNA mediated knockdown of FGFR4. Similarly, FGFR4 knockdown by siRNA (Figure S4F) did not increase sensitivity to estrogen deprivation in BT474, MDA-MB-330, or CAMA1 cells (Figure S4G-I). Consistent with the absence of effects on response to endocrine therapy, FGFR4 overexpression or knockdown alone did not alter pFGFR4 (Y642), pAKT (S473) or pMAPK (T202/Y204) levels (Figure 4G), suggesting that modulating FGFR4 levels alone was not to effect signaling and downstream activity.

Multiple FGF ligands have been reported to be able to activate FGFR4 signaling, including FGF1, 2, 4, 6, 8, 17, 19, 21 and 23 [39, 40]. To explore if ligand stimulation was key to pathway activation and detection of FGFR4 mediated phenotypes, we selected our T47D model with the most robust Dox-inducible overexpression of FGFR4 and assessed the effect of FGFR4 overexpression on downstream signaling in the presence of FGF ligands. As shown in Figure 4H, FGFR4 overexpression alone (PBS group) did not significantly enhance downstream signaling. However, in the presence of FGF1, 2, 4, 6, 8, 17, 19 and 21, FGFR4 overexpression consistently increased pFGFR4 (Y642) and pAKT (S473) levels, and increased pMAPK (T202/Y204) levels in the presence of FGF1, 8 and 17. To see if this ligand stimulation might confer endocrine resistance, we examined the growth of T47D cells in CSS conditions in the presence or absence of FGF ligands and with or without FGFR4 overexpression. Results showed that FGF1, 4, 6 and 8 can significantly promote T47D cell growth in CSS but this was not further enhanced by FGFR4 expression, suggesting that these ligands may be primarily acting via other FGFR family members (Figure 4H). Indeed, FGFR4 overexpression inhibited cell proliferation in the presence of FGF1, 4, 6 and 8, possibly due to competition for FGF ligands between FGFR4 and other FGFRs. FGFR4 overexpression also failed to enhance MDA-MB-134 and MCF7 cells growth in CSS in the presence of FGF ligands (Figure S5). Together, these data suggest that while FGF ligands can promote cell growth in CSS, we have little evidence that this is acting via FGFR4 in the models tested.

To investigate if the previously described FGFR4 R388 SNP [41, 42] was required to mediate resistance to endocrine therapy, we generated Dox inducible FGFR4 (R388) overexpressing T47D cells (Figure S6A). T47D cells showed a robust response to E2 deprivation, but this was not rescued by FGFR4 (R388) overexpression (Figure S6B). We examined the effect of FGFR4 R388 overexpression in the presence of FGF ligands. Similar to FGFR4 (G388) (Figure 4I), FGFR4 (R388) overexpression did not notably promote T47D cell growth in CSS (Figure S6C).

Two FGFR4 hotspot mutations (N535K, V550M) have been identified as enriched in breast cancer metastases [8, 13]. To examine whether the hotspot mutations could enhance FGFR4 signaling and mediate endocrine resistance, we generated Dox inducible FGFR4-N535K and FGFR4-V550M overexpressing MDA-MB-134 cells (Figure S6D). While the overexpression of FGFR4-N535K and FGFR4-V550M increased pFGFR4 (Y642) levels in the absence of FGFs (Figure S6D), they did not alter cellular response to ICI 182780 treatment (Figure S6E, S6F).

### Therapeutic potential of targeting FGFR4 in advanced breast cancer

Considering that FGFR4 is frequently upregulated in metastases from ER+ breast cancer, and the availability of multiple FGFR4 specific inhibitors, we investigated the potential of combination therapy targeting ER and FGFR4 for treating advanced breast cancer. To this end, we treated MDA-MB-134, MCF7 and SUM44PE cells with a combination of ICI 182780 and H3B-6527. All cell lines showed a robust response to ICI 182780, but a weak response to H3B-6527 with no synergy detected with the drug combination (Figure 5A-C).

**Figure 5.**
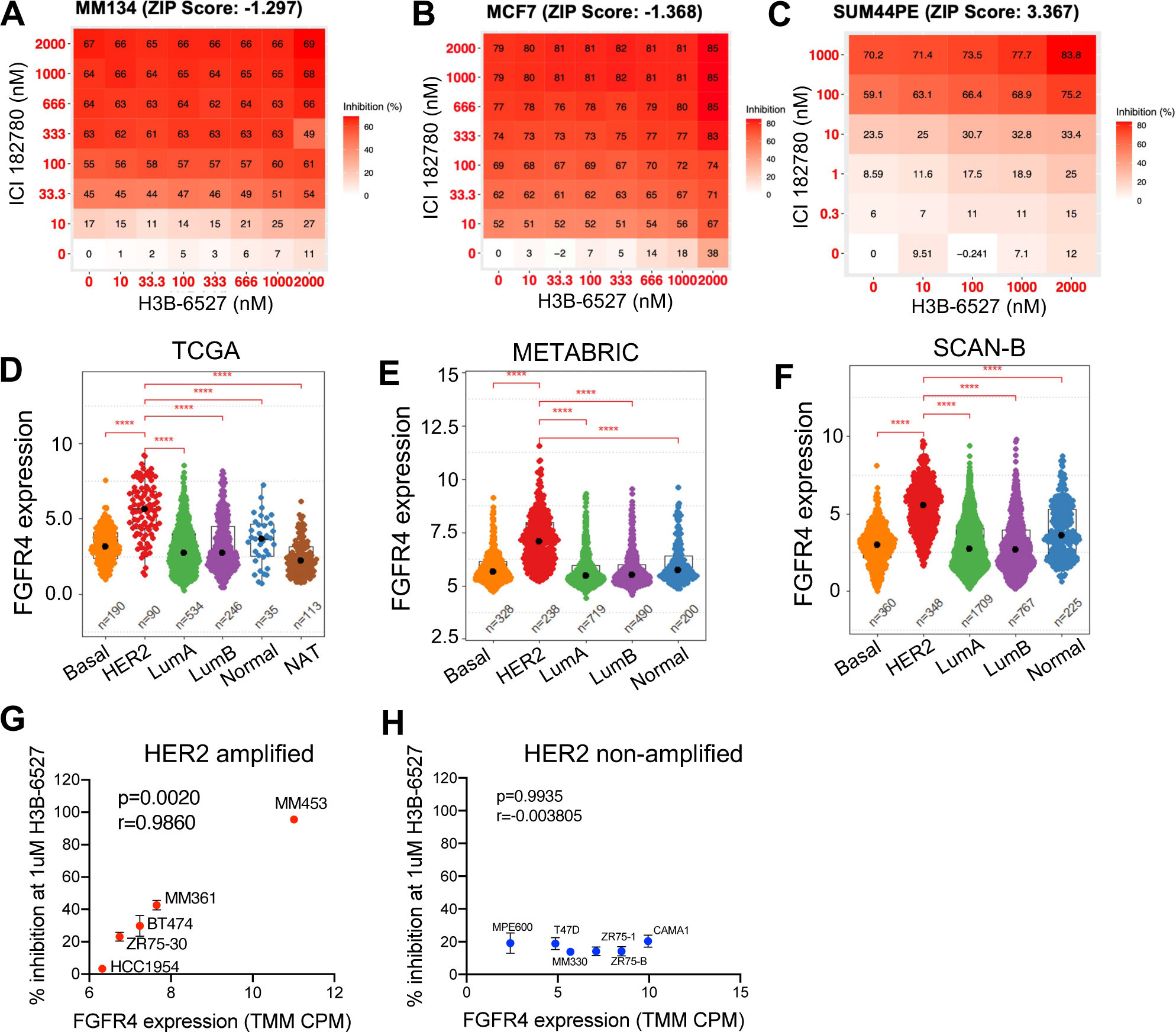
Therapeutic potential of targeting FGFR4 in advanced breast cancer. (A) 2D growth assay of MDA-MB-134, (B) MCF7 and (C) SUM44PE cells in response to ICI 182780 + H3B-6527 combination treatment. Cell number was quantified by FluoReporter assay at day 7. Percentage of cell growth inhibition relative to Veh. treated group was plotted. Synergy score was calculated using zero interaction potency model by R package SynergyFinder. Representative data from N=2 independent repeats for C, N=1 for A and B (D) FGFR4 mRNA expression in TCGA, (E) METABRIC, and (F) SCAN-B data sets across PAM50 subtypes. p values comparing FGFR4 expression in HER2 and other subtypes were calculated by wilcox test. (G) Spearman correlation between FGFR4 expression and percentage of cell growth inhibition in response to 1uM H3B-6527 in HER2 amplified and (H) non-amplified cell lines. Viable cell numbers were quantified using PrestoBlue assay at day 7. FGFR4 expression and HER2 copy number status in cell lines were determined based on CCLE study. r. spearman correlation coefficiency. Error bars represent standard deviation. Representative data from N=2 independent repeats for G and H. * p < 0.05, ** p < 0.01, *** p < 0.001, **** p < 0.0001.

Multiple co-receptors have been reported to interact with FGFR4 and promote cancer progression, including NCAM, N-cadherin, EphA4 and ephexin1, MT1-MMP1, and HER2 [9]. It is plausible that a specific molecular background is required for FGFR4 activation in breast cancer. Therefore, we assessed FGFR4 expression across PAM50 subtypes of breast cancer. Consistent with a previous report [34], FGFR4 showed the highest expression in the HER2 subtype across all data sets analyzed, including TCGA, METABRIC and SCAN-B (Figure 5D-F), while FGFR1-3 showed comparable expression across PAM50 subtypes (Figure S7A). Furthermore, HER2_pY1248 protein levels had the highest positive correlation with FGFR4 expression based on TCGA data (Figure 2A), suggesting that HER2 may play a role in FGFR4 signaling activation. To investigate if the HER2 subtype of breast cancer is more dependent on FGFR4 compared to other subtypes, we examined the response to H3B-6527 in a panel of HER2 amplified and HER2 non-amplified cells based on CCLE study [43] using 2D growth assay (Figure S7B) and colony formation assay (Figure S7C). These studies showed that HER2 amplified cells are more sensitive to FGFR4 inhibition by H3B-6527 than non-amplified cells, which was also strongly correlated with FGFR4 expression (Figure 5G-H), suggesting that high HER2 levels may create a permissive environment for FGFR4 signaling.

## Discussion

While endocrine therapy has revolutionized the treatment of ER+ breast cancer, resistance and metastasis remain a clinical challenge warranting further investigation. We previously reported the overexpression of FGFR4 in 100% (12/12) LTED cell models and in 90% (27/29) of ER+ breast cancer metastases relapsed after endocrine therapy, suggesting a potential role in mediating endocrine resistance [8]. In this study, we demonstrated the upregulation of an FGFR4 signature but not FGFR4 mRNA itself in ER+ primary breast cancer post neoadjuvant endocrine therapy. Further, we found that high FGFR4 expression predicts a worse outcome of patients with ER+ breast cancer. Our data demonstrated that FGFR4 overexpression is at least in part due to ER inhibition by endocrine therapy, since ER represses FGFR4 expression. However, modulation of FGFR4 expression in ER+ breast cancer cell lines did not change their response to endocrine therapy *in vitro*. Specific genomic backgrounds such as HER2 expression seem to be required for FGFR4 functioning in breast cancer.

FGFR4 copy number amplification is rarely observed in breast cancer, yet there is a frequent upregulation in FGFR4 mRNA expression in ER+ breast cancers post endocrine therapy [44, 45]. Through global gene expression correlation analysis, we identified ER as the most significant gene negatively associated with FGFR4 expression. ER is a master transcriptional regulator of ER+ breast cancer that can regulate the expression of genes driving breast cancer progression [46, 47], including growth factor receptors [48] [49]. Here we demonstrate ER as a negative regulator of FGFR4 expression at both the mRNA and protein levels in a panel of ER+ breast cancer cell lines. Such regulation was not observed in MCF7 and SUM44PE cells, suggesting that specific co-regulators might be required for ER mediated regulation of FGFR4 expression. The cross-talk between ER and FGFR signaling including FGFR1 and FGFR2 has been reported to drive endocrine resistance [50–54], but studies are scarce on the role of FGFR4. We hypothesized that the upregulation of FGFR4 as a result of ER inhibitory therapy may mediate resistance to endocrine therapy. Two ER binding sites were identified 63 kb upstream and 22 kb downstream of the FGFR4 transcriptional start site. Further experiments blocking these sites are required to confirm if the regulation of FGFR4 expression is indeed controlled by ER via these regions.

Despite the consistent overexpression of FGFR4 in multiple LTED cells, we did not detect an increased response, in terms of cell growth or colony formation ability, to FGFR4 knockdown or specific inhibitors in LTED cells compared to paired parental cells. This suggests that FGFR4 may not be required for estrogen independent growth of LTED cells, but it does not rule out a role for FGFR4 signaling in the early stages of development of endocrine resistance in these models. To test this hypothesis, we investigated whether the overexpression or knockdown of FGFR4 could alter the response to endocrine therapy at the early stage of treatment using Dox inducible FGFR4 overexpression or knockdown models in a panel of ER+ cell lines. Despite robust overexpression or knockdown of FGFR4, we did not observe a significant growth or survival phenotype change in response to endocrine therapy in a 7-day growth assay or a one-month colony formation assay. Protein analyses showed that FGFR4 overexpression or knockdown alone does not alter pFGFR4 (Y642) levels and downstream signaling activation. We determined that in the presence of FGF ligands, FGFR4 overexpression enhanced pFGFR4 (Y642) and pAKT (S473) levels, suggesting that FGFs are required for pathway activation. Even so, no noticeable increase of cell growth in CSS was observed with FGFR4 overexpression compared to parental cells in the presence of FGF ligands. Interestingly, FGF1/4/6/8 can promote T47D cell growth in CSS, though this was partially decreased following FGFR4 overexpression suggesting that FGFR4 may be competing for ligands with other FGFRs since FGF1/4/6/8 are reported to have a higher affinity to FGFR1-3 than FGFR4 [39, 40]. Further studies knocking down FGFR1-3 are needed to confirm which FGFR is responsible for the FGF mediated increased cell growth in CSS.

FGF19 is considered a specific ligand for FGFR4 with interaction of beta-klotho [40] and other tumor models have suggested this pathway to be a key signaling axis [55, 56]. However, in the presence of FGF19, FGFR4 signaling and the growth of T47D cell were only slightly enhanced following FGFR4 overexpression, suggesting that FGF19 might have a weak potential to activate FGFR4 in T47D breast cancer cells. Further studies exploring the co-expression of beta-klotho or other FGF co-receptors may be required to maximally activate FGFR4 in breast cancer.

Multiple co-receptors have been reported to interact with FGFR4 to promote tumor progression suggesting that a specific genetic background may be required for FGFR4 signaling [13]. Consistent with Garcia-Recio S et al. [34], our data showed that FGFR4 is expressed at the highest level in the HER2 subtype of breast cancer, while FGFR1-3 displays comparable expression across PAM50 subtypes in TCGA, METABRIC and SCAN-B data sets. In addition, HER2 (Y1248) expression is the most highly correlated protein with FGFR4 in TCGA data. Our studies also show that HER2 amplified cell lines demonstrate a stronger response to the FGFR4 specific inhibitor H3B-6527 than HER2 non-amplified cells with the magnitude of response also correlated with FGFR4 expression. Together, these finding support a potential role for HER2 in activation of FGFR4 signaling in breast cancer. Garcia-Recio S et al. demonstrated a role for FGFR4 in regulating the luminal to HER2 enriched subtype switching, further suggesting an interplay between FGFR4 and HER2 signaling [34]. A recent study also identified FGFR4 as an essential gene following anti-HER2 treatment and showed synergistic effect of anti-FGFR4 with anti-HER2 therapy in breast cancer [57].

Although FGFR4 and a FGFR4 signature are frequently upregulated post endocrine therapy and high FGFR4 expression predicts poorer survival of patients with ER+ disease, our experimental data reported here does not support a general role of FGFR4 overexpression or mutations in mediating endocrine resistance in ER+ breast cancer cell lines *in vitro.* However, we note that our study has limitations. The cell models we explored likely have varied expression of different FGFR4 co-regulators and this should be explored to comprehensively investigate the role of FGFR4 in mediating cell growth and treatment resistance in different genetic backgrounds. Secondly, although we did not observe a role for FGFR4 in mediating endocrine resistance *in vitro*, it is possible that *in vivo* findings may be different. The tumor microenvironment, especially FGFs secreted by fibroblasts, likely plays an important role in FGFR4 activation.

In summary, our data suggests that despite upregulation in endocrine resistant tumors and cell models, FGFR4 might not be an effective drug target for ER+ advanced breast cancer, however the HER2 amplified subtype of breast cancer may benefit from strategies targeting FGFR4 [19] which should be further explored, emphasizing the importance of precision cancer treatment.

## Methods

### FGFR4 gene signature analysis

RNA sequencing data of ER+ primary breast cancer was downloaded from GSE59515 + GSE5537 [36] and dbGaP (phs000472) [35]. Data was normalized to log2 trimmed mean counts per million for GSE59515 + GSE5537 and log2 fragments per kilobase of transcript per million mapped reads for dbGaP (phs000472). A FGFR4 activation gene signature was generated by Cejalvo JM et al. [34]. Genes with false discovery rate < 1, fold change < 0.5 and not included in cluster 4 proliferative related genes are kept as FGFR4 induced genes and listed below. A GSVA [58] method was used to calculate FGFR4 induced gene signature score based on normalized data. FGFR4 induced genes: RGS16, DAPL1, CYP2F1, FCGR2B, CNTNAP2, ANXA1, MGAT3, TMEM45A, SLPI, EFEMP1, CXCL1, SLC39A8, ABCC11, PXDNL, MEI1, EPAS1, PHLDA1, MET, MYH6, SLC5A1, SLC6A14, HSPB8, GULP1, RAG1, DNAH8, PRKCA. Paired Wilcoxon test was used to calculate statistical difference between samples pre and post endocrine therapy.

### Survival Analysis

The ER+ invasive lobular carcinoma (ILC) samples were previously collected at the Pittsburgh Biospecimen Core and UPMC Magee-Women’s Hospital. Briefly, 123 ER+ ILC samples were collected from patients over the age of 18 with a diagnosis of primary ILC. Subjects were excluded for a previous history of breast cancer, metastasis at diagnosis, concurrent non-breast tumor at diagnosis, or ER-disease. Subjects were censored at date of diagnosis of second primary, including contralateral breast cancer, date of death unrelated to breast cancer or at date of last follow-up in the absence of any other qualifying event. Samples were subjected to Nanostring quantification of 695 genes expression, including FGFR4. Raw counts were collected from the nCounter Digital Analyzer and transferred to the nSolverTM software (v 2.5) for data analysis. Raw counts were normalized to 5 invariant reference probes (count normalization). Normalized FGFR4 expression value and patient clinical and distant recurrence free survival information is provided in supplementary table 1. Molecular Taxonomy of Breast Cancer International Consortium (METABRIC) data was downloaded from the Synapse software platform (syn1688369; Sage Bionetworks, Seattle, WA, USA) [59], and patients with ER+ disease were included for the survival analysis. For both cohorts, patients were stratified based on FGFR4 expression, with top third quantile defined as FGFR4 high and the rest defined as FGFR4 low. Log-rank test was performed to examine the statistical difference in survival between patients with high and low FGFR4 expression.

### FGFR4 correlation analysis

Gene expression data from The Cancer Genome Atlas (TCGA) and Sweden Cancerome Analysis Network-Breast (SCAN-B) project, and TCGA RPPA level 4 processed data were downloaded from the Gene expression Omnibus database [GEO: GSE62944], Gene expression Omnibus database [GEO: GSE96058], and Cbioportal (https://www.tcpaportal.org/tcpa/download.html) respectively. METABRIC data was downloaded as described above. RNAseq counts from TCGA and SCAN-B study were normalized to log2 counts per million (CPM) using EdgeR package [60]. Microarray data from METABRIC was processed as described in previous publication [59]. Spearman correlation analysis was performed between FGFR4 mRNA and TCGA RPPA protein expression as well as FGFR4 mRNA and ESR1 mRNA in TCGA, METABRIC and SCAN-B cohorts.

### Cell culture and estrogen deprivation

Cell lines used in this study are mainly purchased from ATCC, including MDA-MB-134 (also abbreviated as MM134), MCF7, MDA-MB-361 (MM361), HCC1954, MDA-MB-453 (MM453), CAMA-1, T47D, BT-474, MDA-MB-330 (MM330), ZR-75-1, and ZR-75-30. SUM44PE was purchased from Asterand. BCK4 cells were obtained as a gift from Dr. Britta Jacobsen developed as detailed [61].

Culture media for each cell line used in this study are listed below: MM134: 1:1 DMEM: L-15 with 10% fetal bovine serum (FBS; Life Technologies), MM134-LTED-B/ MM134-LTED-E/SUM44PE-LTED-A/SUM44PE-LTED-B: IMEM with 10% charcoal stripped serum (CSS, Life Technologies), SUM44PE: DMEM/F12 with 2% CSS and additional supplements as previously described [62], BCK4: MEM with nonessential amino acids (Life Technologies) and insulin (Sigma-Aldrich) and 10% FBS, MCF7/MM361/HCC1954/MM453/CAMA-1: DMEM with 10% FBS, T47D/BT-474/MM330/ZR-75-1/ZR-75-30/ZR-75-B/MPE600: RPMI with 10% FBS. All culture media were purchased from Thermo Fisher Scientific. For estrogen deprivation, cell culture media was changed to IMEM + 10% CSS and refreshed 2-3 times a day for a total of 3 days.

All cell lines were maintained in 5% carbon dioxide and cultured with Falcon tissue culture dish or flask (fisher scientific). Cell lines were split and culture media were refreshed twice a week. Cell lines were cultured for less than 6 months at a time, routinely tested to be mycoplasma free.

### Generation of doxycycline inducible FGFR4 overexpression and knockdown models

Wild type and mutant FGFR4 genes were cloned into pInducer20 plasmid (addgene #44012), a third generation lentivirus vector. pInducer20 with FGFR4 gene inserted was then transfected into HEK293T cells with third generation lentivirus packing vectors pMDL, Rev, and VSVG using polyethylenimine (Sigma-Aldrich # 408727). Virus was harvested from HEK293T cell culture supernatant at 24 and 36 hours post transfection and used to infect target cells twice with the adding of polybrene (v/v, 1:1000) (EMD Millipore # TR-1003-G). 24 hours post the second infection, virus containing media was removed from the target cells, and fresh virus free media was added to cells to let cell recover from infection stress. 24 hours later, 1mg/ml G418 (Thermo Fisher Scientific # 10-131-027) was added to target cells to select for cells successfully infected. G418 was refreshed every week until uninfected control cells are all dead. For dox inducible FGFR4 knockdown models, FGFR4 shRNA and Renilla control shRNA were cloned into LT3GEPIR plasmid (addgene #111177), a third generation lentivirus vector. The same protocol was applied for target cell infection and selection. 1 ug/ml puromycin (Life Technologies, A11138-03) was used for selecting infected cells.

### siRNA transfection

ON-TARGETplus SMARTpool small interfering RNAs (siRNAs) were purchased from Dharmacon: FGFR4 (#L-003134-00-0005) and ESR1 (#L-003401-00-0005). Cells were reversed transfected with 1 pmol (96-well) or 25 pmol (6-well) siRNA using Lipofectamine RNAiMAX (Life Technologies, 13778150) and Opti-MEM (Thermo Fisher Scientific) following manufacturer’s protocol. RNA and proteins were harvested 48 hours post transfection.

### RNA extraction and quantitative PCR (qPCR)

RNA was extracted from cells using RNeasy mini kit (Qiagen #74106) according to the manufacturer’s protocol. RNA concentration and quality were measured by NanoDrop. mRNA was reversed transcribed into cDNA using PrimeScript™ RT Master Mix (Takara Bio #RR036B). cDNA was then used for qPCR with SsoAdvanced Universal SYBR (Bio-Rad #1726275) using FGFR4 primers: forward: TGCAGAATCTCACCTTGATTACA / reverse: GGGGTAACTGTGCCTATTCG and RPLP0 primers: forward: GAGTATGATCCTACCAGACCCTTC / reverse: CCTGATCATGGAGGGTCAAATC. FGFR4 expression of each sample was normalized to RPLR0 internal control to adjust for the variation of cDNA loaded.

### Western blot (WB)

Protein was extracted from cells using RIPA buffer (50mM Tris pH 7.4, 150mM NaCl, 1mM EDTA, 0.5% Nonidet P-40, 0.5% NaDeoxycholate, 0.1% SDS) supplied with 1x protease and phosphatase inhibitor (Thermo Fisher #78442). Cells were incubated in RIPA buffer on ice for 15 min followed by a sonication at 100 AMP for 10 min (30s on and 30s off) to further break down the cells. Protein lysates were separated from cell debris by centrifugation at 13,000 rpm for 15 min at 4 degree and quantified using a BCA assay (Thermo Fisher #23225) according to manufacturer’s protocol. 40 ug protein per sample was loaded into and separated by a 10% SDS-PAGE gel, and then transferred to a PVDF membrane (EMD Millipore # IPFL00010). The membrane was then blocked in Intercept blocking buffer (LiCOr #927-40000) for one hour at room temperature, and then probed with primary antibody at 4 degree overnight. Primary antibody information is listed below: FGFR4 (Cell Signaling #8562, 1:1,000), ER-Alpha (Abcam #131105, 1:1,000), pFGFR4(Y642) (signalway #11836-2, 1:1,000), pFRS2(Y196) (Cell Signaling #3864s, 1:1,000), pAkt (S473) (Cell Signaling #9271S. 1:1,000), Pp44/42MAPK(T202/Y204) (Cell Signaling #4377, 1:1,000), and Beta-Actin (Sigma A5441, 1:1,000).

PBST (PBS + 0.1% Tween-20) was used to wash membrane three times before probing the membrane with fluorophore labelled secondary antibody for one hour at room temperature (anti-mouse 680LT: LiCor #925-68020, anti-rabbit 800CW: LiCor #925-32211, 1:10,000). Membrane was then washed by PBST and imaged by LiCOr Odyssey CLx Imaging system.

### FGFR4 signaling study

0.5 to 1 million cells were plated into 6-well plate and treated with 0.5 or 1 mg/ml Dox (Sigma-Aldrich #D9891) to induce FGFR4 overexpression or knockdown for 48 hours. Cells were then starved in serum free media overnight, and treated with 10% serum, PBS plus 1 ug/ml heparin (Sigma-Aldrich # H3149), 20 mg/ml FGF1/2/4/6/8/17 plus 1ug/ml heparin, or 100ng/ml FGF19/21/23 for 15 min. FGF1 was from R&D Systems. FGF2 was from Fisher Scientific. All the other FGFs were from PeproTech. The expression of FGFR4 and its downstream signaling mediators were measured by WB as described above. Antibody information was listed in WB method.

### Cell growth and colony formation assay

For cell growth assay, 5,000 cells for IDC/NST cell lines and 15,000 cell for ILC cell lines were seeded into each 96-well. Vehicle or serial dilution of treatments were then added to cells, including Dox, Tamoxifen (Tam, Selleckchem #s1238), Fulvestrant (ICI 182780) (Selleckchem #S1191), BLU554, H3B-6527 and FGFs. BLU554 and H3B-6527 are provided by Blueprint Medicines and H3 Biomedicine respectively. Specific treatment conditions for each experiment are described in corresponding figure legends. For growth assay in estrogen deprived condition, cells were maintained in IMEM + 10% CSS media. Cell number was quantified at day 7 post treatment using FluoReporter (Thermo Fisher Scientific #F2962) or PrestoBlue (Thermo Fisher Scientific # A13262) assay according to the manufacturer’s protocol.

For colony formation assay, 8,000 cells for MDA-MB-134, SUM44PE, MDA-MB-361, ZR75-30, ZR75-1, CAMA1, BT474, MDA-MB-330 and BCK4, and 3,000 cells for T47D, MCF7, ZR75-B, HCC1954, MDA-MB-453 and MPE600 were seeded into each well of a 12-well plate. Vehicle or drugs were then added at the time of cell seeding and refreshed at day 7. Cell plates were collected 14 days post initial treatment, fixed in ice-cold 100% methanol on ice for 10 min, then stained with 0.5% crystal violet (Sigma-Aldrich, #C0775) solution for 10 min at room temperature. Extra dye was washed off with water. Stained crystal violet was dissolved in 10% glacier acetic acid and read for absorbance at 560 nm to quantify relative colony numbers. For one-month colony formation assay in CSS, 250,000 MM134 and 50,000 MCF7 cells were seeded in each well of a 12-well plate. Culture media, Dox, E2 and Tam were refreshed every week according to each treatment condition specified in corresponding figure legend for 4 weeks. Cell culture plates were scanned by Incucyte live imaging system and the area of well covered by cells was quantified by the default soft wear installed with Incucyte.

## Acknowledgments

We would like to thank The Cancer Genome Atlas (TCGA) Research Network (https://www.cancer.gov/tcga), the Sweden Cancerome Analysis Network, and the Molecular Taxonomy of Breast Cancer International Consortium (METABRIC) for generating and sharing their data to empower our study. We appreciate technical support from Jian Chen and Loise Mazur, and support from personnel the Department of Pathology at UPMC Magee-Womens Hospital, and the Pittsburgh Biospecimen Core (PBC). This research was supported in part by the University of Pittsburgh Center for Research Computing, RRID:SCR_022735, through the resources provided. Specifically, this work used the HTC cluster, which is supported by NIH award number S10OD028483. The funders had no role in study design, data collection and analysis, decision to publish, or preparation of the manuscript.

## Ethics declarations

Conflict of interest

The authors have no conflicts of interest to disclose.

## Ethical approval

All clinical samples were collected with informed consent and approval from the Institutional Review Board (IRB) at the University of Pittsburgh

---

**Supplementary figure 1.**
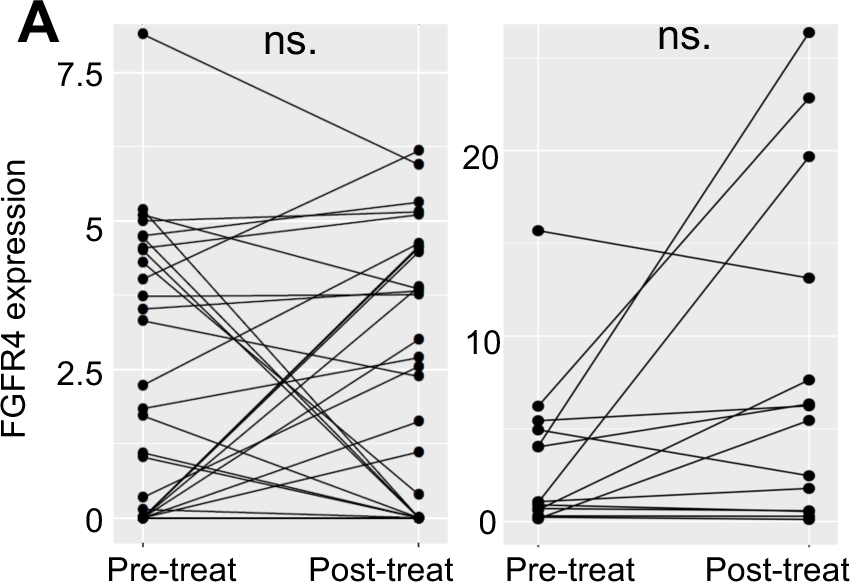
FGFR4 expression pre- and post-neoadjuvant aromatase inhibitor treatment in ER+ primary breast cancer. (A) FGFR4 expression pre-and post-90 days of neoadjuvant aromatase inhibitor treatment in ER+ primary breast cancer using data from GSE59515+GSE5537 (left) and dbGaP (phs000472) (right). p value was calculated by paired Wilcoxon test. ns. (nonsignificant)

**Supplementary figure 2.**
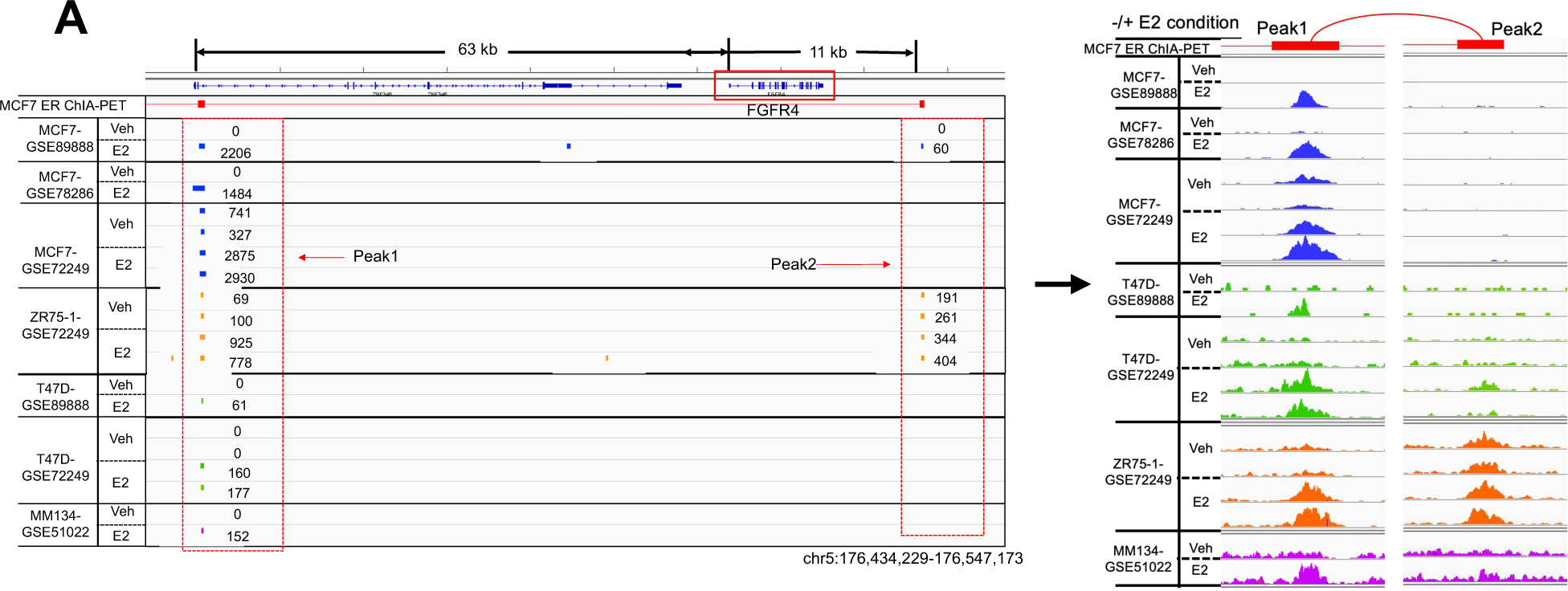
ER binding peaks around FGFR4 gene based on ChIP sequencing. (A) ER binding peaks around FGFR4 gene identified from ChIP sequencing data of MCF7, ZR75-1, T47D and MM134 cells treated with vehicle or estradiol. GSE number of source data were as noted in the figure.

**Supplementary figure 3.**
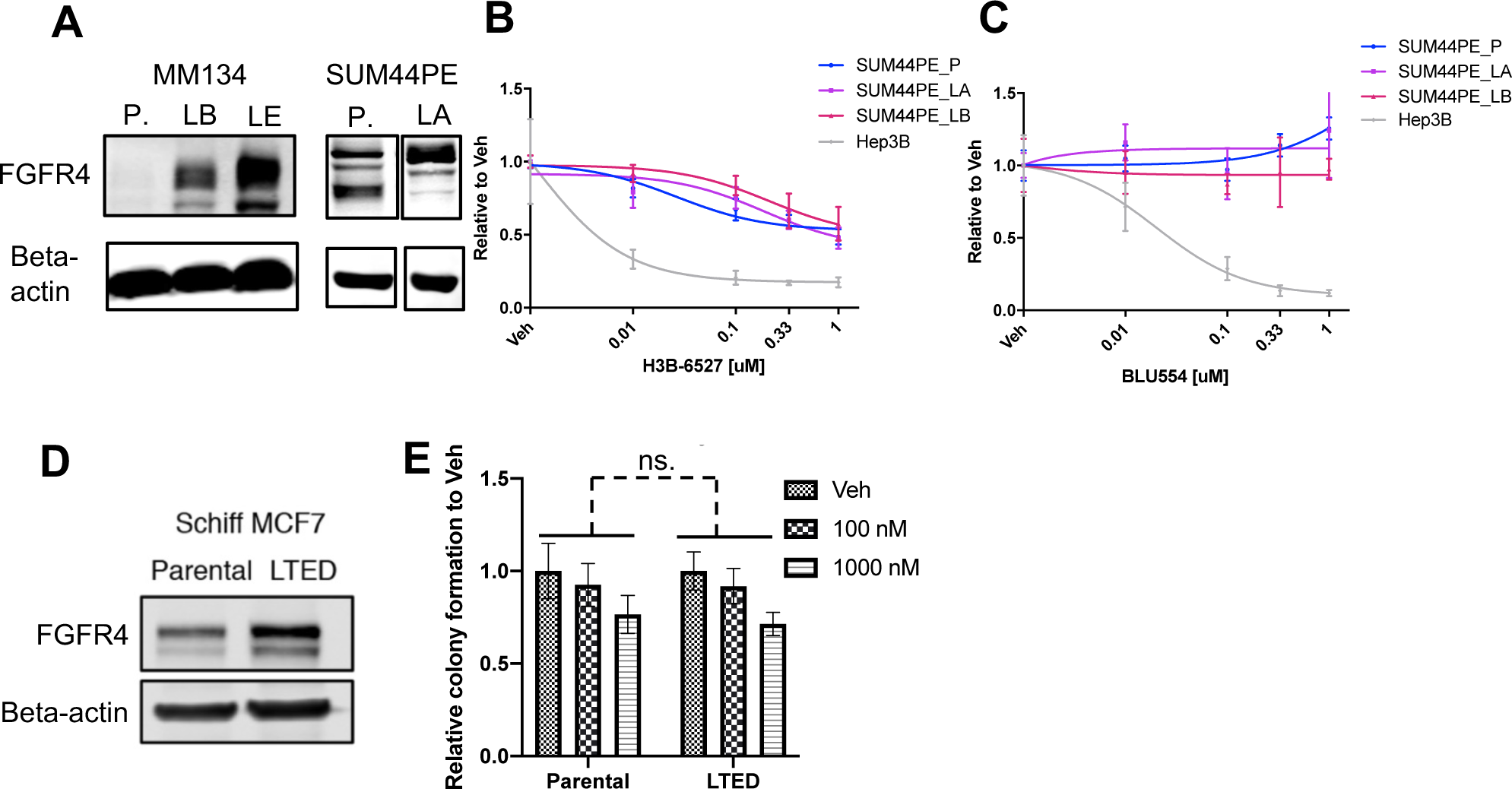
FGFR4 expression and response to FGFR4 inhibition in LTED cells and paired parental cells. (A) Expression of FGFR4 in MDA-MB-134 and SUM44PE LTED cells and paired parental cells quantified by western blot. N=1 experiment. (B) SUM44PE parental (P), LTED-A (LA) and LTED-B (LB) cells dose response to H3B-6527 and (C) BLU554 measured by 2D growth assay. Cell number was quantified by FluoReporter assay at day 7. Data was normalized to Veh. treated group. Curves were simulated using nonlinear regression model with variable slopes by PRISM. Hep3B was used as a positive control for FGFR4 inhibitors. N=1 experiment. (D) FGFR4 expression in MCF7 parental and LTED cells quantified by western blot. N=1 experiment. (E) MCF7 parental and LTED cells response to H3B-6527 treatment measured by 2D colony formation assay for 14 days. Colonies were quantified by crystal violet staining. Data was normalized to Veh. treated group. p value was calculated by two-way ANOVA. Representative data from N=2 independent repeats Error bars in all figures represent standard deviation. ns. (nonsignificant)

**Supplementary figure 4.**
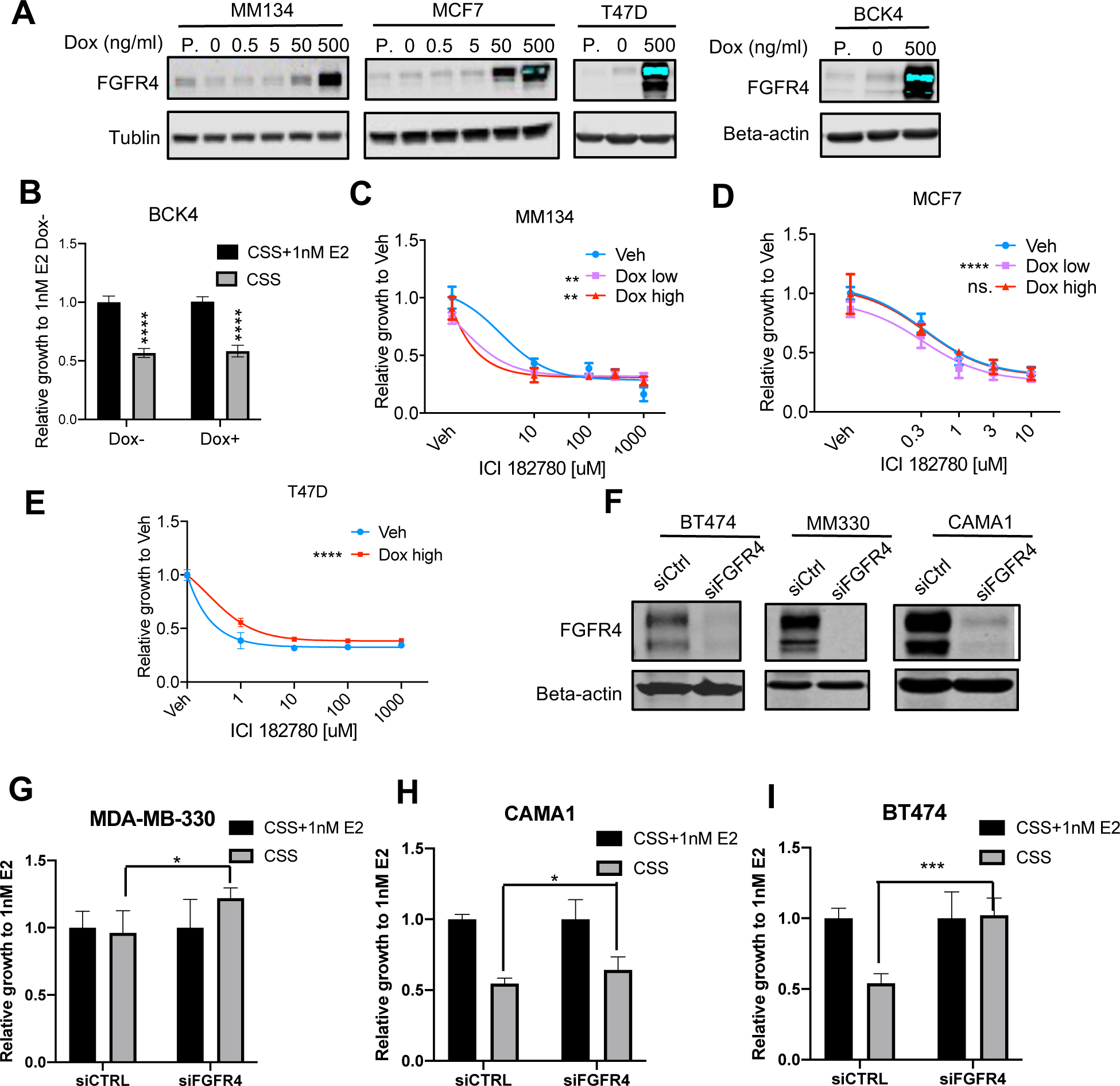
FGFR4 expression alterations cannot change response to endocrine therapy in cell models tested. (A) Dox induced FGFR4 overexpression in MDA-MB-134, MCF7, T47D, and BCK4 cells measured by western blot. Representative data from N=3 independent repeats. P. parental cells. (B) Relative growth of BCK4 in CSS supplied with or without E2 and treated with or without Dox induced FGFR4 overexpression. Viable cell number was quantified by PrestoBlue at day 7. Data was normalized to +E2 -Dox condition. N=1 experiment. (C) MDA-MB-134, (D) MCF7, and (E) T47D response to ICI 182780 with or without Dox induced overexpression of FGFR4. Cell number was quantified by FluoReporter assay at day 7. Data was normalized to Veh. treated group of each condition. Dose response curve was simulated using nonlinear regression model with variable slopes by PRISM. N=1 experiment (F) FGFR4 protein expression in BT474, MDA-MB-330, and CAMA1 cells with or without siFGFR4 medicated knockdown of FGFR4, quantified by western blot. N=1 experiment. (G) Relative growth of BT474, (H) MDA-MB-330, and (I) CAMA1 in CSS supplied with or without 1 nM E2 and treated with or without siFGFR4. Data was normalized to +E2 group of each condition. N=1 experiment All error bars represent standard deviation. p values for B and G-I were calculated by t test, and for C-E were calculated by two-way ANOVA. * p < 0.05, ** p < 0.01, *** p < 0.001, **** p < 0.0001. ns. (nonsignificant).

**Supplementary figure 5.**
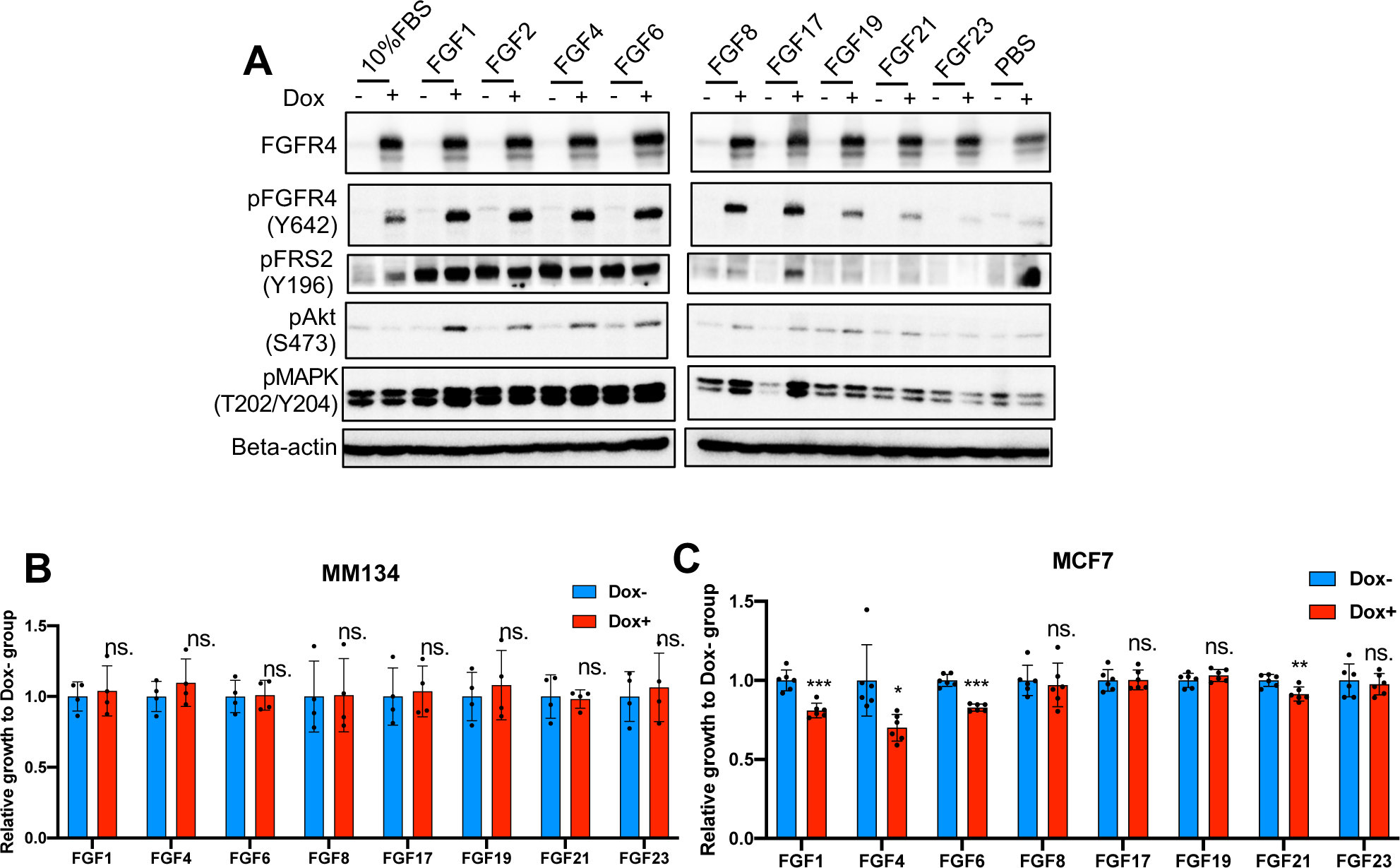
FGFR4 signaling is enhanced following FGFR4 overexpression in the presence of FGF ligands, which however, cannot promote cell growth in CSS. (A) Western blot assessment of FGFR4 and downstream signaling in T47D cells. Cells were induced with Dox for 2 days, serum starved overnight, then stimulated with FGFs (20 ng/ml for FGF1/2/4/6/8/17, 100ng/ml for FGF17/19/21/23) for 15 min and harvest for protein. N=1 experiment. (B) Relative growth of MDA-MB-134 and (C) MCF7 cells in CSS with or without Dox induced FGFR4 overexpression in the presence or absence of FGFs (20 ng/ml for FGF1/4/6/8/17, 100ng/ml for FGF17/19/21/23). Viable cell number was quantified by PrestoBlue assay at day 7. Data was normalized to Dox-group of each treatment condition. p values comparing Dox+ and Dox-group was calculated. N=1 experiment All error bars represent standard deviation. p values comparing Dox-and Dox+ group for B, and C were calculated by t test. * p < 0.05, ** p < 0.01, *** p < 0.001, **** p < 0.0001. ns. (nonsignificant).

**Supplementary figure 6.**
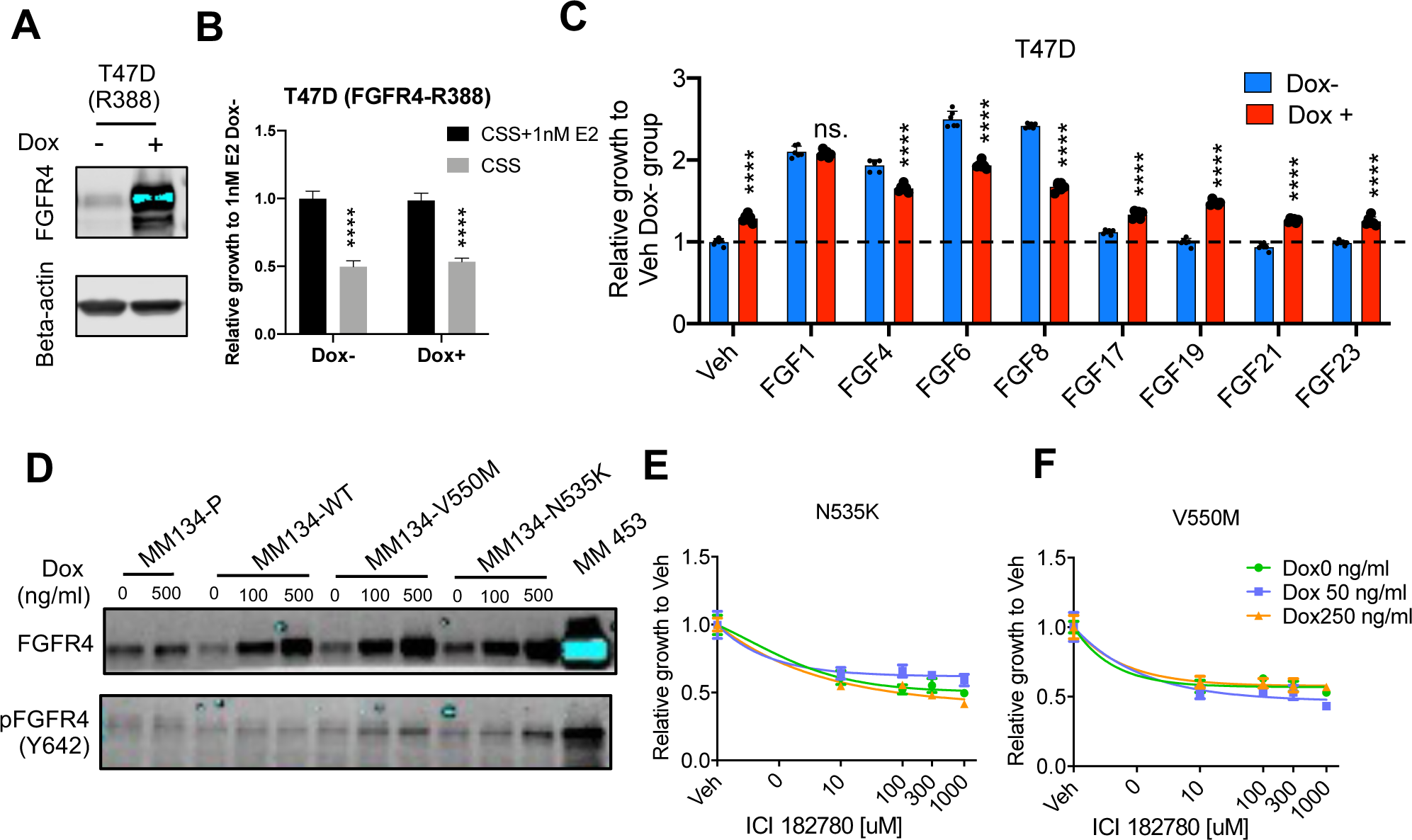
FGFR4 G388R single nucleotide polymorphism and hotspot mutations cannot mediate endocrine resistance. (A) WB of FGFR4 in T47D cells with or without Dox induced FGFR4 (R388) overexpression. (B) Relative growth of T47D cells in CSS with or without E2 with or without FGFR4 (R388) overexpression. Viable cell number was quantified by PrestoBlue assay at day 7. Data was normalized to CSS+1nM E2 Dox-group. N=1 experiment (C) Relative growth of T47D cells in CSS with or without Dox induced FGFR4 (R388) overexpression in the presence or absence of FGFs (20 ng/ml for FGF1/4/6/8/17, 100ng/ml for FGF17/19/21/23). Viable cell number was quantified by PrestoBlue assay at day 7. Data was normalized to Dox-Veh. treated group. N=1 experiment (D) FGFR4 and pFGFR4 (Y642) levels in MM134 cells with or without Dox induced overexpression of WT (wild type) and mutant (V550M, N535K) FGFR4 quantified by western blot. N=1 experiment. (E) MDA-MB-134 response to ICI 182780 with or without Dox induced overexpression of FGFR4-N535K and (F) FGFR4-V550M. Cell number was quantified by FluoReporter assay at day 7. Data was normalized to Veh. treated group. Dose response curve was simulated using nonlinear regression model with variable slopes. N=1 experiment. All error bars represent standard deviation. p values for B and C were calculated using t test. * p < 0.05, ** p < 0.01, *** p < 0.001, **** p < 0.0001.

**Supplementary figure 7.**
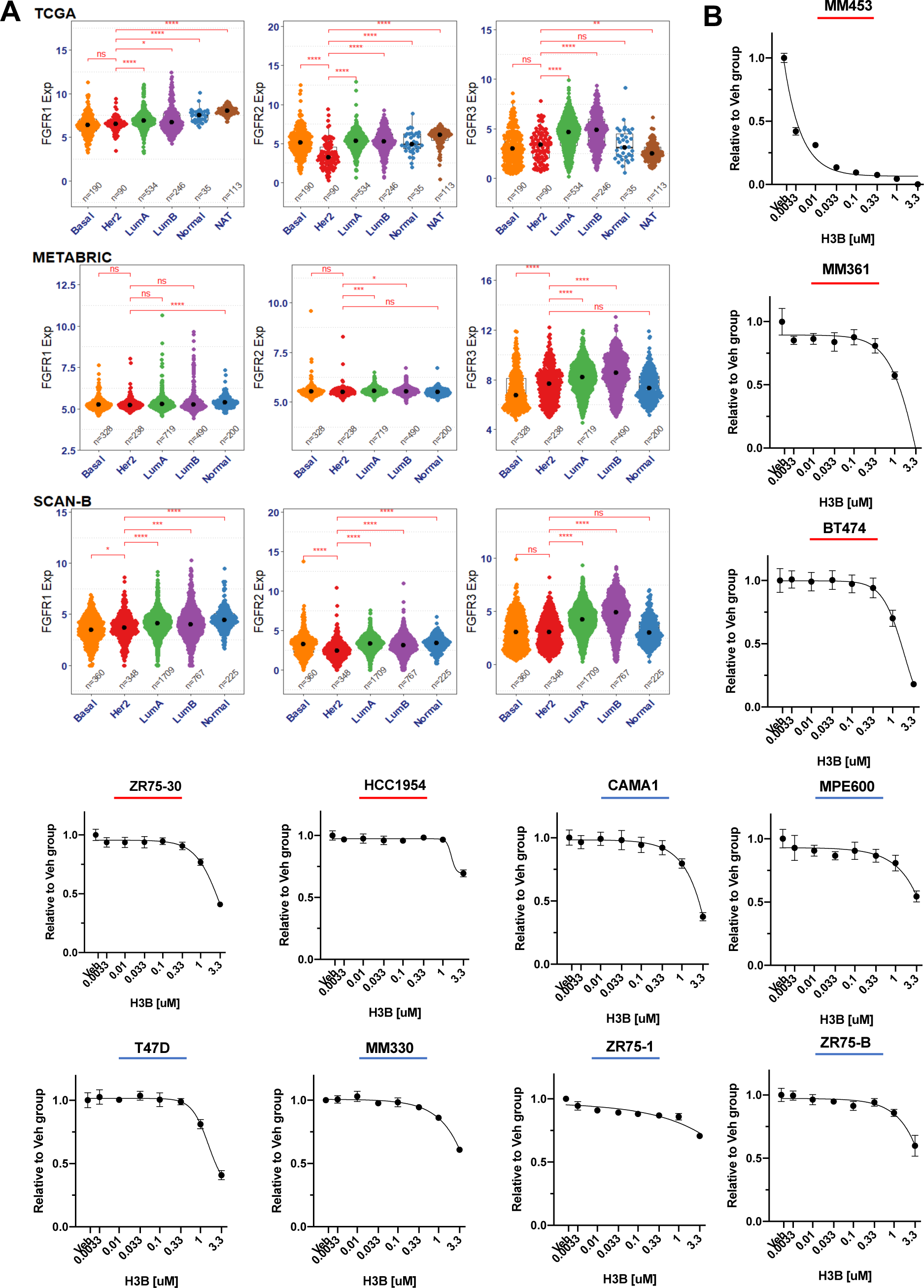
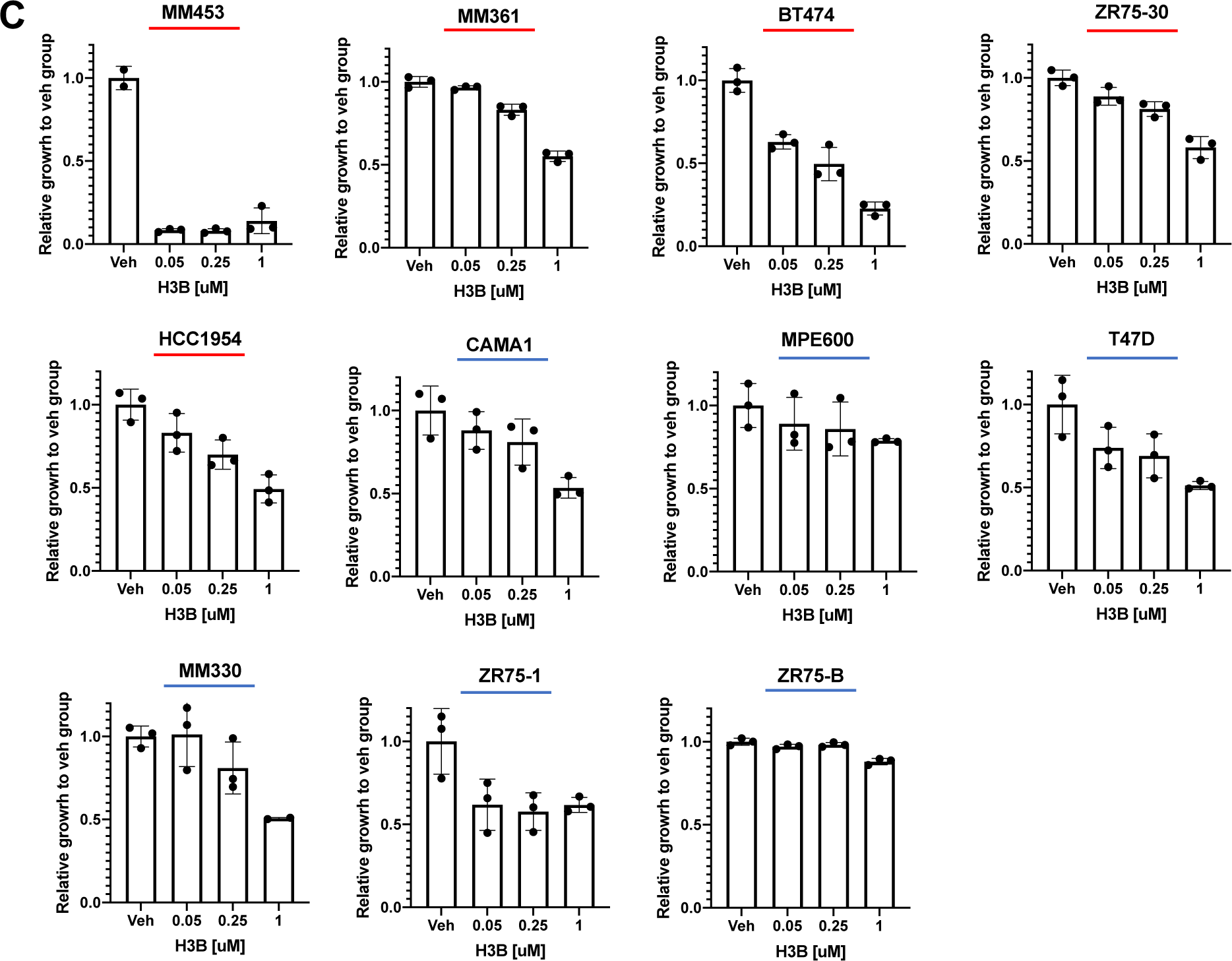
FGFR1-3 expression across PAM50 subtypes and the response to H3B-6527 in HER2 amplified and HER2 non-amplified cells. (A) Expression of FGFR1-3 in TCGA, METABRIC, and SCAN-B datasets across PAM50 subtypes. p values comparing FGFR1-3 expression in HER2 versus other subtypes were calculated by Wilcox test. * p < 0.05, ** p < 0.01, *** p < 0.001, **** p < 0.0001. (B) Response to H3B-6527 in HER2 amplified (red underline) and non-amplified (blue underline) cell lines measured by 2D growth assay. Viable cell number was quantified by PrestoBlue assay at day 7. Data was normalized to Veh. treated group. Dose response curve was simulated using nonlinear regression model with variable slopes by PRISM. Representative data from N=2 independent repeats. (C) Response to H3B-6527 in HER2 amplified (red underline) and non-amplified (blue underline) cell lines measured by colony formation assay at day 14. Cell number was quantified by crystal violet staining. Data was normalized to Veh. treated group. N=1 experiment.

